# PAREameters: computational inference of plant microRNA-mRNA targeting rules using RNA sequencing data

**DOI:** 10.1101/710814

**Authors:** Joshua Thody, Vincent Moulton, Irina Mohorianu

## Abstract

MicroRNAs (miRNAs) are short, non-coding RNAs that influence the translation-rate of mRNAs by directing the RNA-induced silencing complex to sequence-specific targets. In plants, this typically results in cleavage and subsequent degradation of the mRNA. This can be captured on a high-throughput scale using degradome sequencing, which supports miRNA target prediction by aligning degradation fragments to reference mRNAs enabling the identification of causal miRNA(s). The current criteria used for target prediction were inferred on experimentally validated *A. thaliana* interactions, which were adapted to fit that specific subset of miRNA interactions. In addition, the miRNA pathway in other organisms may have acquired specific changes, e.g. lineage-specific miRNAs or new miRNA-mRNA interactions, thus previous criteria may not be optimal. We present a new tool, PAREameters, for inferring targeting criteria from RNA sequencing datasets; the stability of inferred criteria under subsampling and the effect of input-size are discussed. We first evaluate its performance using experimentally validated miRNA-mRNA interactions in multiple *A. thaliana* datasets, including conserved and species-specific miRNAs. We then perform comprehensive analyses on the differences in flower miRNA-mRNA interactions in several non-model organisms and quantify the observed variations. PAREameters highlights an increase in sensitivity on most tested datasets when data-inferred criteria are used.

## INTRODUCTION

Small RNAs (sRNAs) are short, non-coding RNAs with roles in transcriptional and post-transcriptional gene regulation in eukaryotes [1]. In plants, the latter mode of action is achieved via a class of sRNAs, the microRNAs (miRNAs), which reduce the amount of messenger RNA (mRNA) available for translation by directing the RNA-induced silencing complex (RISC) to their sequence-specific mRNA target(s) and inducing cleavage and subsequent degradation of the mRNA [2]. The Parallel Analysis of RNA ends (PARE) protocol [3], also known as degradome sequencing, captures the 5’ ends of down-stream cleaved mRNAs, which are used to quantitatively predict miRNA-mRNA interactions.

Improvements to next-generation sequencing technologies have resulted in larger and more diverse experiments, including multi-omics ones [4]. It has also led to the sequencing and annotation of many different organisms’ genomes and facilitated functional studies outside of the context of model organisms [5]. However, a vast proportion of our understanding of specific biological mechanisms is based on the study of model organisms, mostly due to their lower regulatory complexity and availability of extensive, varied public sequencing datasets. Many computational methods designed for extracting information/features from sequencing data, (e.g. sRNA classification and target prediction) often summarize the data-mining results into rule-based models, derived from experimental observations. However, this approach carries the risk of overfitting a model (e.g. set of thresholds or accepted ranges) on particular species or specific set of observations. Recently, this was acknowledged for miRNA classification criteria [6, 7], which were recently updated [8].

Current target prediction tools use fixed criteria that were inferred on low-throughput, validated *A. thaliana* miRNA-mRNA interactions [9, 10]. Subsequent analyses, in various organisms, have shown that these criteria do not capture all known and expressed miRNA-mRNA interactions (in *A. thaliana* [11], and *O. sativa* [12]). In addition, the portability of these criteria between various organisms and tissues has not been yet quantitatively evaluated. Recent tools for degradome assisted sRNA target prediction, including PAREsnip2 [13] and sPARTA [14], were used to assess the suitability of the *Allen et al*. criteria [9]. Results on PARE datasets against a set of experimentally validated interactions in *A. thaliana* [13] revealed that only ~80% of the expressed and experimentally validated interactions were reported when using the *Allen et al*. criteria [9]. Further analyses revealed that the remaining ~20% were missed mostly due to discrepancies in the number or position of mismatches, gaps, G:U pairs and the minimum free energy (MFE) ratio. This suggests that the current criteria may be too stringent or over-fitted on a small set of organism specific, experimentally validated miRNA-mRNA interactions. Moreover, the sensitivity and precision of the prediction may differ based on the size or characteristics of the input data. For example, the functional analysis of a specific miRNA may benefit from reduced precision, yet good sensitivity, to increase the number of candidates for further investigations; whereas, an analysis on the entire set of sRNAs requires simultaneously high sensitivity and precision.

To illustrate the issues with the current fixed criteria, we further assessed the variation in accuracy of different sets of targeting rules. Using an *A. thaliana* leaf dataset (D1), we employed two sets of targeting criteria, the *Allen et al*. rules [9] and the criteria we manually inferred from a set of validated *A. thaliana* miRNA-mRNA interactions [13] (ST1)). These criteria were provided as input parameters for PAREsnip2 [13], which is a highly-configurable tool for the prediction of sRNA targets from sequencing data (paired sRNA and degradome). Evaluation of these criteria showed an increase in sensitivity from 78.5-81.4% to 94.5-96.2%, with precision values of 88.7-92.1% to 82.1-85.9% for the *Allen et al.* criteria and the manually inferred criteria, respectively (ST2), over three biological replicates. Upon further inspection, the majority of validated interactions that were missed using the manually inferred criteria were due to having an MFE ratio less than the selected cut-off value of 0.65.

In this paper, we describe a new tool, PAREameters, which leverages the flexibility of PAREsnip2 targeting parameters. Using publicly available datasets, we illustrate how PAREameters facilitates the inference of targeting criteria based on information extracted from paired sRNA and degradome data in conjunction with miRNA annotations (e.g. from miRBase v22 [15]). The tool is freely available, open source and provided as part of the UEA sRNA Workbench [16].

## MATERIALS AND METHODS

### The PAREameters pipeline

In Figure 1 we give an overview of the PAREameters pipeline. It is applied to synonymous sRNA and PARE samples to infer miRNA targeting rules. The input to PAREameters consists of sRNA and corresponding degradome samples (technical or biological replicates can be used for assessing technical variation and noise between samples or for the exclusion of spurious results). An annotated reference genome and transcriptome, and a set of known plant miRNAs (e.g. from miRBase [15]) are also required. The PAREameters tool was implemented in Java (version 8); the code used to create the plots and perform the significance tests is implemented in R (version 3.5.1, Apple Darwin) and can be invoked from the PAREameters pipeline using system calls, assuming a valid version of R is installed and correctly configured. All computational analyses and benchmarking were performed on a desktop machine running Ubuntu 18.04 equipped with a 3.40GHz Intel Core i7-6800K six core CPU and 128GB RAM.

**Figure 1.**
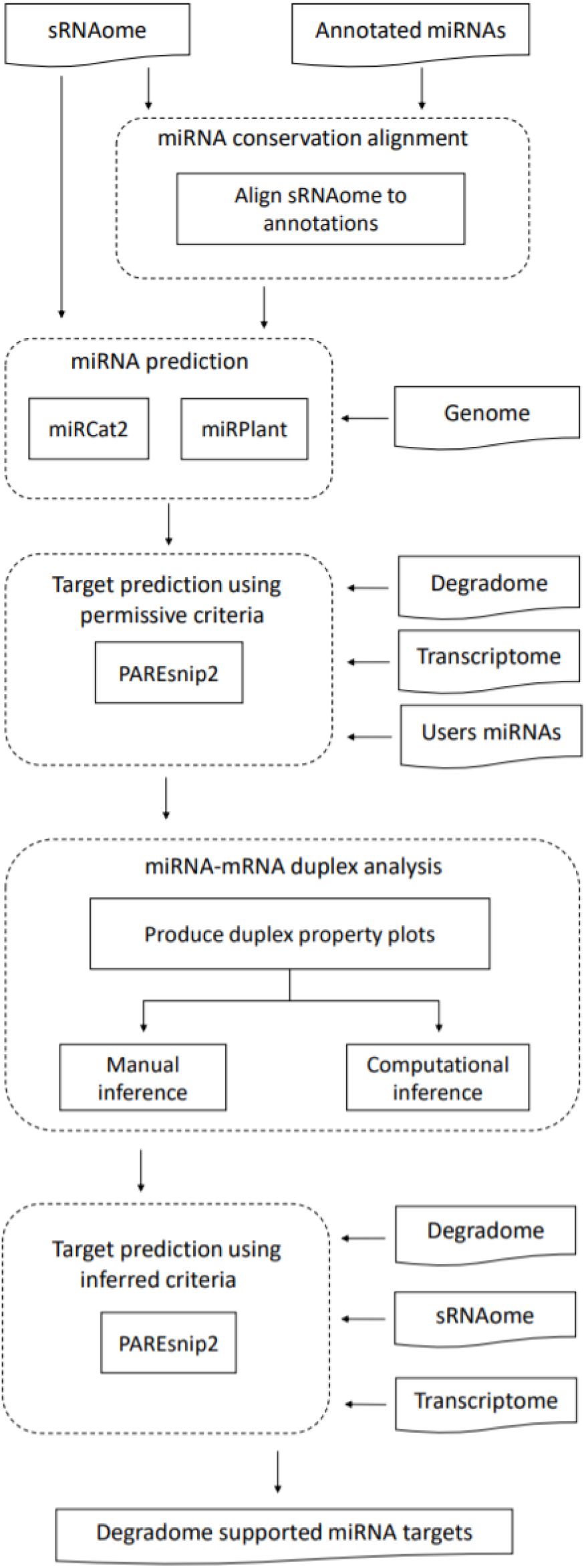
PAREameters pipeline. The input and output data are represented by curved rectangles; the processes are represented by straight rectangles. PAREameters takes as input two types of sequencing samples, paired sRNA and degradome, a genome with corresponding annotations and current miRBase miRNA annotations. The output is a set of data-inferred thresholds for a rule-based prediction of miRNA-mRNA interactions using e.g. PAREsnip2.

To remove low quality reads, sequencing errors or to identify sample outliers, PAREameters includes several optional filtering methods: (i) sequences containing ambiguous bases (e.g. Ns) are discarded; (ii) a low sequence complexity filter is applied based on the single, di- or tri-nucleotide frequencies [13], with set thresholds of 75%, 37.5% and 25%, respectively; (iii) if a reference genome is provided, all reads that do not align to it are discarded.

### Selection of the miRNA candidates

The miRNA candidates used as input for the PAREameters pipeline can be obtained via two approaches: (a) with focus on conserved miRNAs. The input sRNA samples are aligned (+/+ strand only) to all known plant miRNA sequences, obtained from miRBase [15], allowing up to 2 mismatches and no gaps. The selected sequences are then used as input to miRPlant [17]. Candidates that fulfil the criteria for miRNA prediction (default tool criteria), are then retained for the subsequent steps; (b) with focus on all miRNAs (conserved and new) as predicted using miRCat2 [18] with the whole sRNA sample as input. In addition, to compensate the stringent criteria of miRNA prediction tools, the user can provide their own annotated miRNA entries if they have an abundance ≥5 (user-defined parameter) but did not fulfil the criteria of the prediction tools.

### Permissive target prediction

Small RNAs that satisfy miRNA biogenesis criteria (as described above) are provided as input to PAREsnip2 [13]. The target prediction is then performed on the input data using a set of highly-permissive, user-configurable, parameters (ST1).

The miRNA-mRNA interactions predicted by PAREsnip2 are kept if the abundance of the peak of interest is ≥5 and are further classified into high-confidence (HC) or low-confidence (LC). For the former the peak is the highest across the whole transcript (i.e. category 0 or 1); for the latter the peak is not the highest on the transcript (i.e. category 2 or 3). Peaks with abundance <5 are excluded because it is difficult to distinguish between true miRNA cleavage products and random degradation at such low abundance.

### Duplex analysis and inference of targeting criteria

Valid miRNA-mRNA duplexes, based on the analysis of the degradome data coupled with the miRNA prediction, are characterized using specific properties, such as the number and location of mismatches, G:U wobble pairs and adjacent mismatches, the alignment score and the minimum free energy (MFE) ratio. The algorithm then infers a set of targeting criteria that attempts to retain at least 85% (user-defined parameter) of the valid miRNA-mRNA duplexes. We chose the default value of the retain rate parameter based on the optimal variation of the sensitivity and precision values of inferred criteria across an incremental range of retain rate values on a benchmark leaf *A. thaliana* dataset comprising of 3 replicates [13], presented in the results. The biological interpretation of the retain rate threshold is that a higher degree of complementarity between a miRNA and its target results in higher confidence that the interaction is genuine, whereas interactions with weaker complementarity may require further experimental validation.

Using a set of experimentally validated interactions as validation, we focused on HC interaction pairs at known target sites with corresponding miRNAs [13]. The validation classes, true positives (TP), false positives (FP) and positives (P) are used in a loose sense i.e. TP consists of the predicted interactions with experimental validation, FP is the set of predicted interaction for which, currently, there is no experimental validation, and P is the set of experimentally validated interactions. For each set of targeting rules, we present the sensitivity as Se = TP/P (number of predicted validated interactions) and the precision as PPV = TP / (TP+FP) (proportion of additional predicted interactions, which are not currently validated).

In addition, as the PAREameters pipeline provides a summary of the interaction properties to the user, manual curation of the results allows the user to choose a set of targeting criteria that satisfies their choice of sensitivity and precision. The value suggested as default for the retain rate parameter corresponds to the smallest overall value of the absolute ratio between Se and PPV since it corresponds to the minimum increase of sensitivity with respect to the loss in PPV (ST3).

The significance of the distribution of properties with respect to the miRNA was calculated using offset χ^2^ tests and the contribution of each property was assessed using individual Fisher exact tests [19]. To better understand the contribution of specific nucleotide base-pairs or motifs, the distributions of match pairs (A/U, G/C and G/U), mismatch pairs (A/m, C/m, G/m, U/m) and gap pairs (A/g, C/g, G/g, U/g) were also determined, with the positional significance evaluated using the χ^2^ test and the individual Fisher exact tests. Lastly, the distribution of miRNA-mRNA duplex MFE ratios [9, 13] were analysed using Kolmogorov-Smirnov tests.

### Datasets

Three *A. thaliana* datasets were used, for each set paired sRNA and PARE samples were available: (D1) wild-type leaf triplicates (D1A, D1B and D1C), GSE90771 (sRNAs) [18] and GSE113958 (PARE) [13]; (D2) wild-type leaves in a growth time-series at 35 days (D2A), 45 days (D2B) and 50 days (D2C), GSE55151 [20]; (D3) wild-type flower (D3A), leaf (D3B), root (D3C) and seedling (D3D) of plants grown at 15°C, NCBI BioProject PRJNA407271 [21]. The genome and transcriptome versions are TAIR10 and both were obtained from The Arabidopsis Information Resource [22]. The set of experimentally validated *A. thaliana* miRNA/mRNA interactions were obtained from a previous study [13].

In addition to the *A. thaliana* datasets, we exemplify the usage of PAREameters on sRNA and corresponding PARE datasets from *A. trichopoda* leaf (D4A) and opened female flower (D4B) (GSE41811); *G. max* leaf (D5) (GSE76636) [23]; *O. sativa* inflorescence (D6) (GSE18251) [24] and *T. aestivum* 2.2mm spikes (D7) (GSE36867) [25]. The transcriptome and genome sequences for organisms other than *A. thaliana* were obtained from EnsemblPlants Release 43 [26], namely, *A. trichopoda* genome version AMTR1.0, annotation version AMTR1.0, *G. max* genome version 2.1, annotation version 2.1, *O. sativa* genome version IRGSP-1.0, annotation version IRGSP-1.0, *T. aestivum* genome version IWGSC (genome build accession GCA_900519105.1), annotation version IWGSC.

Summaries about each dataset, such as the number of raw and unique reads and genome matched reads can be found in ST4. In addition, for the sRNA data, we report the number of known miRNAs present (based on current miRBase [15] annotation) and for the PARE data, we also include the number of transcriptome matching reads.

## RESULTS

### Evaluation the inferred targeting rules in *A. thaliana*

First we compare the increase in accuracy of the computationally inferred targeting rules versus the manually inferred *Allen et al.* rules. Next, we assess the stability of the inferred rules using random subsampling of the input data. The effect of additional parameters, such as size of the input data and the user-defined retain rate, are also discussed.

When comparing the sensitivity of the manually inferred criteria and the *Allen et al*. rules on the set of *A. thaliana* validated interactions, the former outperformed the latter by ~15%. However, this increase in performance may be due to the over-fitting of the targeting criteria on the currently known interactions. In addition, due to the scarcity of validated interactions (either as number of valid interactions or localization of specific modes of action in different cell types [27]), these criteria may not be portable between various organisms or tissues. Therefore, we used the PAREameters tool to infer targeting criteria from the *A. thaliana* D1, D2 and D3 datasets. The inferred criteria were then utilized by PAREsnip2 for target prediction and the results evaluated and compared to the *Allen et al.* rules. The resulting parameters reported by the PAREameters pipeline can be found in ST5. The evaluation method used is identical to that of the manually inferred criteria. Specifically, for each dataset, the class of positive (P) data included experimentally validated miRNA-mRNA interactions with HC transcript peaks and corresponding miRNA sequence with abundance ≥5.

The results, presented in Table 1, show that the computationally inferred criteria provide increased sensitivity compared to the *Allen et al*. criteria, whilst also maintaining precision on most datasets. Over all datasets, PAREameters inferred criteria with a median sensitivity of 88.5% (range: 82.8-89.4%) vs 81.4% (range: 75.6-84.6%) for the *Allen et al.* criteria. The median precision for the inferred criteria was 91.3% (range: 80.1-96.8%) vs 91.4% (range: 83.8-97.5%) for the *Allen et al*. criteria.

**Table 1:** Comparison of sensitivity and specificity between the *Allen et al*. criteria and the PAREameters inferred criteria on the *A. thaliana* datasets. The latter lead to an increased sensitivity; the apparent loss in precision may be due to the incomplete characterization (and validations) of regulatory interactions and can only increase as more experimentally confirmed interactions become available.

| Dataset | # miRNAs | # V | Allen V | Inferred V | Allen NV | Inferred NV | Allen Se | Inferred Se | Allen PPV | Inferred PPV | Se gain | PPV difference |
| --- | --- | --- | --- | --- | --- | --- | --- | --- | --- | --- | --- | --- |
| D1A | 37 | 129 | 105 | 112 | 9 | 8 | 81.395% | 86.822% | 92.105% | 93.333% | 5.426% | 1.228% |
| D1B | 38 | 131 | 109 | 116 | 11 | 10 | 83.206% | 88.550% | 90.833% | 92.063% | 5.344% | 1.230% |
| D1C | 35 | 121 | 95 | 107 | 12 | 14 | 78.512% | 88.430% | 88.785% | 88.430% | 9.917% | -0.355% |
| D2A | 40 | 140 | 117 | 125 | 14 | 29 | 83.571% | 89.286% | 89.313% | 81.169% | 5.714% | -8.144% |
| D2B | 38 | 137 | 113 | 121 | 13 | 30 | 82.482% | 88.321% | 89.683% | 80.132% | 5.839% | -9.550% |
| D2C | 40 | 144 | 117 | 120 | 3 | 4 | 81.250% | 83.333% | 97.500% | 96.774% | 2.083% | -0.726% |
| D3A | 32 | 79 | 64 | 68 | 4 | 7 | 81.013% | 86.076% | 94.118% | 90.667% | 5.063% | -3.451% |
| D3B | 29 | 70 | 57 | 58 | 11 | 13 | 81.429% | 82.857% | 83.824% | 81.690% | 1.429% | -2.133% |
| D3C | 36 | 111 | 84 | 98 | 6 | 7 | 75.676% | 88.288% | 93.333% | 93.333% | 12.613% | 0.000% |
| D3D | 35 | 104 | 88 | 93 | 3 | 3 | 84.615% | 89.423% | 96.703% | 96.875% | 4.808% | 0.172% |

We also evaluated the time and memory performance of PAREameters on each dataset. The runtime of the pipeline depends on the size of the input data (sequencing depth of the sRNA and PARE samples and the size of the reference genome). On *A. thaliana* D1, D2 and D3 datasets, the runtime range was 16 minutes and 52 seconds to 1 hour 4 minutes (this excludes the time taken to build the bowtie index as this is only done once per species) and the memory usage varied between 5GB and 8GB (ST6). The inference component of PAREameters is linear on the size of the sRNA and PARE input data.

Based on the properties of HC miRNA-mRNA duplexes with cleavage signal confirmation in the PARE data, PAREameters extracted targeting criteria that increased the sensitivity and retained precision versus existing fixed criteria when tested against a golden standard – the set of experimentally validated interactions in *A. thaliana*. To avoid the overfitting of targeting criteria based on characteristics of the input data, we tested the stability of the inferred properties using a cross-validation technique and the set of experimentally validated *A. thaliana* miRNA-mRNA interactions on the D1, D2 and D3 datasets. Specifically, we used the HC interactions with corresponding miRNA sequences in each dataset as a starting point. We then randomly split the HC validated interactions in each dataset to form two groups: the training group, containing 75% of the data, and the testing group, which contained the remaining 25%. PAREameters was used to infer parameters on the training set and these were employed by PAREsnip2 for target prediction on the test set. We then calculated the sensitivity and precision of the inferred parameters on the training set and on the test set. The random cross-validation was repeated 50x and the results, displayed in ST7, show that PAREameters is able to infer targeting parameters with a median sensitivity of 77%, [range: 66-81%] and precision 83% [range: 75-100%] when evaluated on the unobserved testing data.

The decrease in sensitivity from our previous analysis likely originates from the fact we are inferring criteria from one set of miRNA-mRNA interactions and testing on a different set of miRNA-mRNA interactions. Whereas previously, we were inferring criteria from the whole set of PAREameters predicted HC miRNA-mRNA interactions. This further supports our hypothesis that miRNAs may have different modes of action or target complementarity requirements and demonstrates that using just one set of fixed criteria is not sufficient when performing miRNA target prediction.

To investigate how increasing the amount of training data may lead to a more accurate representation of inferred targeting criteria, we evaluated the computationally inferred criteria produced by PAREameters on different sized subsets of the experimentally validated interactions contained within the D1 datasets. Starting with 10% of the validated data, followed by increments of 10% until the final value of 90%, we used PAREameters to infer criteria on the data subset and then evaluated those criteria on the remaining unseen data. Analysis on each subset was performed 50 times and the results shown in ST8. On each dataset, increasing the amount of training data resulted in an overall increase in sensitivity. Intriguingly, the increase in training data resulted in a decrease in precision. However, this should not be seen as a negative result, as we’ve previously stated, the class FP is the set of predicted interaction for which, currently, there is no experimental validation. Indeed, the current class of positive data is almost certainly incomplete, therefore further experimental validation can only increase the sensitivity and precision values for the inferred criteria.

To evaluate how changes to the PAREameters retain rate parameter impacts Se and PPV, we evaluated the computational inferred targeting criteria produced by PAREameters on the D1 dataset with increasing retain rate values. The results of this analysis are shown in ST9 and SF1. Starting with an initial value of 0.5 and with increments of 0.05 thereafter, we recorded the number of validated and non-validated interactions being captured at each value. Next, we computed the ratio between the Se and the PPV; the data-inferred threshold for the retain rate parameter is the value that minimizes this ratio i.e. it is the value for which the increase in Se is minimal with respect to the loss in PPV (ST3). In the *A. thaliana* D1 data used to exemplify the retain rate the global minimum was also the last local minimum; for samples with multiple local minimum, the default value would indicate the global minimum, however we suggest a visual assessment of the Se/PPV ratio and if local minima have close values then higher values are preferred.

Using the initial value on the D1A dataset, we capture a total of 30 miRNA-mRNA interactions, all of which are experimentally validated interactions. At the other end of the scale, using a retain rate of 1.0 captured 156 interactions, which comprised of 128 validated and 28 non-validated. The default parameter value (0.85) captures a total of 120 interactions and provides a sensitivity value of 86.8% and precision value of 93.3%. A visual representation of these results of all three replicates in D1, which show similar results, can be found in SF1. The threshold of 0.85 minimizes simultaneously the gain on sensitivity and the loss on precision. The 0.85 value was consistent across technical replicates. In experiments for which the values vary between samples, we recommend the usage of a consistent threshold across all samples of the experiment.

### Consistency of attribute distributions and inferred criteria across miRNA subsets in *A. thaliana*

To evaluate the portability of targeting criteria (and distribution of properties) across miRNA subsets we used PAREameters on a set of conserved and species-specific *A. thaliana* miRNAs [15], with the corresponding interactions. The group built on the conserved miRNAs comprised 201 miRNA-mRNA interactions from 42 unique miRNA sequences (ST10). The group built on miRNAs specific to the *Brassicaceae* family comprised 184 interactions from 47 unique miRNA sequences (ST11). The summaries of the position-specific property distributions, which include the localizations of gaps, mismatches and G:U wobbles, and the MFE ratio distributions for the conserved and specific miRNA interactions are presented in Figure 2A and 2B, respectively. In Figure 2A, the *Brassicaceae* specific miRNAs show highly similar results to that of *Allen et al.* [9], e.g. a large proportion of mismatches or G:U wobble pairs at position 1, no mismatches at the canonical positions 9 and 10 and relatively few mismatches in the 5’ core region (positions 2-13) of the miRNA when compared to the 3’ end. In contrast, the requirements for complementary of species-specific miRNAs (Figure 2A) appears to differ considerably when compared to conserved miRNAs, especially at the miRNA 5’ end, with mismatches being tolerated at positions 5, 8 and 9, in addition to the canonical position 10 of the miRNA. To evaluate whether the differences in properties between specific-specific and conserved miRNA interactions in *A. thaliana* are significant, we performed χ^2^ tests of significance using the conserved properties as the expected distribution and the species-specific properties as the observed distribution. Additionally, we use the Fisher’s exact test to determine the specific property at each position responsible for the significance of the differences. The results of the significance analysis for the position-specific property distributions are presented in Table 2. Based on the χ^2^ tests, significance differences (*p*-value ≤ 0.05) between properties can be found at positions 1, 5, 8, 14, 16 and 21. Based on the Fisher’s exact test, position 16 has significant differences in both proportions of mismatches and G:U pairs, positions 5, 8, 14, 20 and 21 have significant differences in proportion of mismatches and positions 1 and 13 have significant differences in the proportion of G:U pairs. We also analysed the differences in MFE ratio distributions between conserved and species-specific miRNAs, shown in Figure 2B, and the significance of the differences were evaluated using the Kolmogorov-Smirnov test, which reported a *p*-value of 8.57×10^−10^. These results may suggest a higher complementarity requirement between conserved miRNAs and their targets than that of species-specific miRNAs.

**Table 2:**
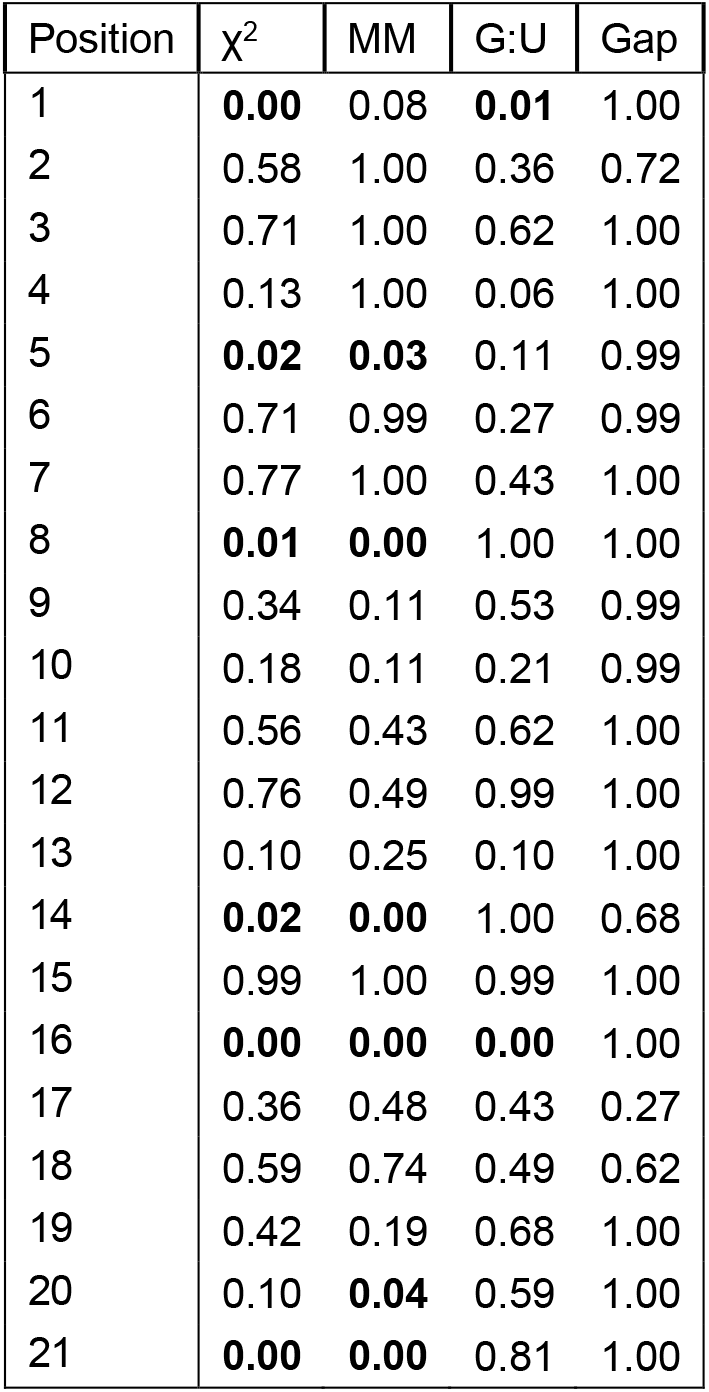
χ^2^ and Fisher’s exact test significance results on the position-specific properties for conserved and species-specific miRNA-mRNA interactions in *A. thaliana*. The contribution of specific properties (such as MM, Gaps and G:U is assessed using Fisher exact tests). Values before the significance threshold (*p*-value ≤ 0.05) are highlighted in bold. The first position and the 8-10nt region (which is essential for inducing cleavage) are either significant or marginally significant, indicating a potential divergence in the type of interactions of either conserved or species-specific miRNAs.

**Figure 2:**
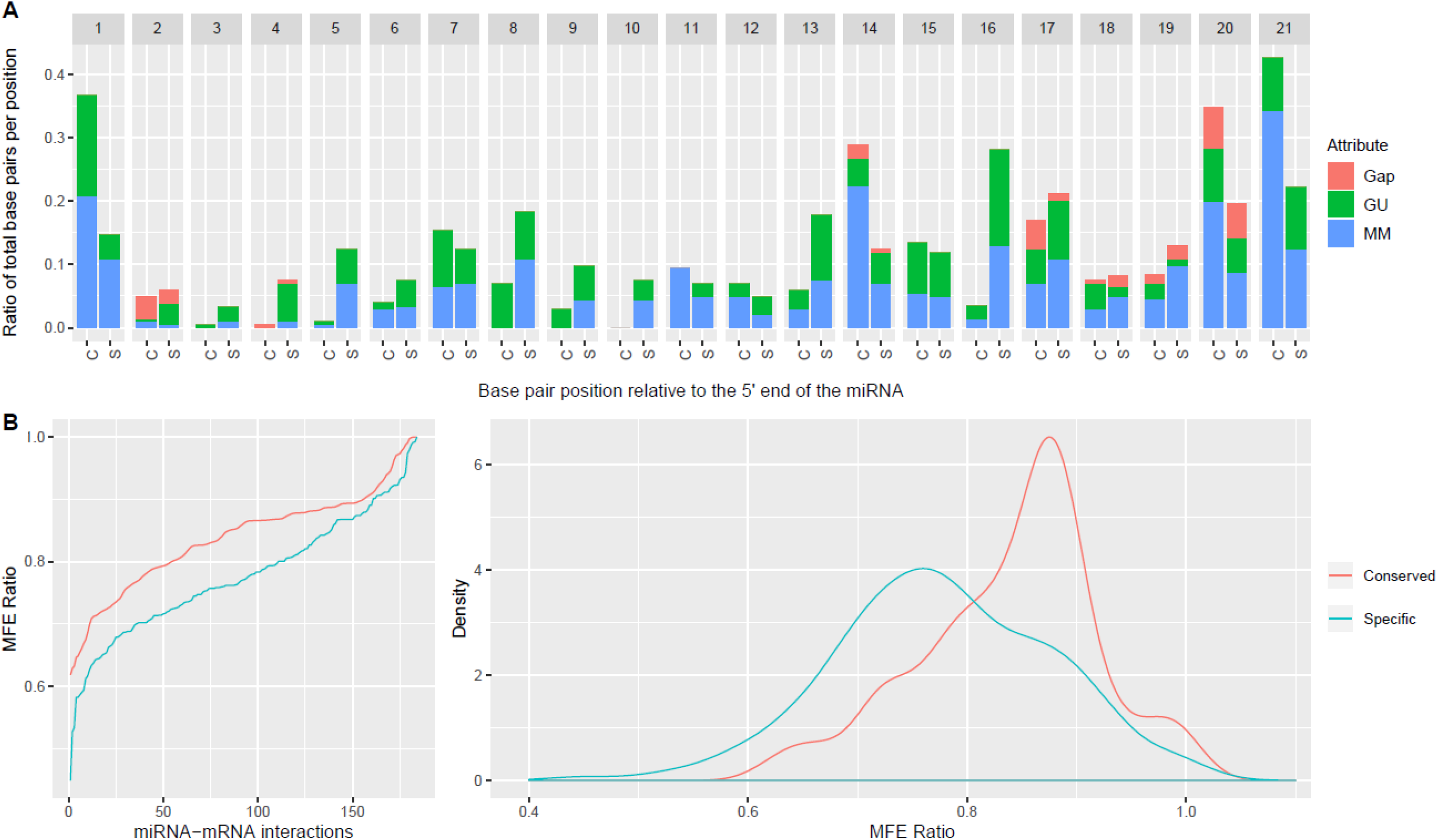
Side-by-side comparison of property distributions for conserved and species-specific miRNAs in *A thaliana*. Using experimentally validated miRNA-mRNA interactions as input, we calculated the position-specific properties (panel A) and the MFE ratio distribution (panel B) for the conserved and species-specific miRNA-mRNA interactions; the former are presented as proportions out of all interactions, in each category and the latter as a cumulative distribution. The significance of the differences in the localization of gaps, G:U pairs and mismatches was assessed using offset χ^2^ tests and the contribution of individual categories was evaluated using Fisher exact tests. The first position and the 8-10 range, important for the cleavage ability of the miRNA, showed significant/marginal significant differences; in addition, positions 14 and 16 illustrated the divergence in properties between these subsets. The similarities in the distributions of the MFE ratios were evaluated using the Kolmogorov-Smirnov test, which reported a *p*-value of 8.57×10^−10^, the distributions of MFE ratios were different both in location of the mode and the shape of the distirbutions.

To investigate the portability between criteria inferred exclusively on conserved or species-specific miRNA interactions, we evaluated the inferred rules of each set of interactions (all four pairwise combinations: conserved rules on conserved interactions, conserved rules on species-specific interactions and the similar pairs on the species-specific rules), using PAREsnip2. The results, presented in Table 3, show a consistent decrease in sensitivity for both the conserved and species-specific miRNAs when inferring criteria on the other subset of miRNA-mRNA interactions. Specifically, a decrease from 82.08% to 65.67% and 76.09% to 55.98% for the conserved and species-specific miRNA-mRNA interactions, respectively. Further investigation into these differences support our previous observation regarding the differences in MFE ratio of conserved and species-specific miRNA interactions, with the inferred values being 0.75 and 0.68, respectively, further supporting our previous observation regarding an increased complementarity requirement for conserved miRNAs. Another intriguing difference between the inferred criteria is an allowed mismatch or G:U pair at position 10 of the species-specific miRNAs.

**Table 3:**
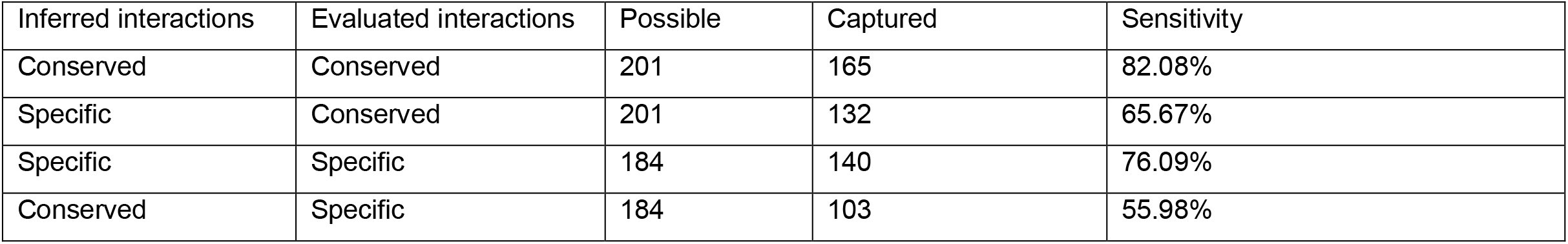
Sensitivity on cross pairwise comparisons for criteria inferred on conserved or species-specific miRNAs for the validated *A. thaliana* interactions. The targeting criteria were inferred using a retain rate of 0.85; a considerable decrease in sensitivity was observed for the mismatched pairs i.e. training on conserved and testing on specific and the symmetric pair, highlighting the impact of data-inferred targeting criteria.

The differences between the properties of conserved and species-specific miRNA-mRNA interactions highlight the need for customization in the set of criteria used for describing and capturing miRNA-mRNA interactions when conserved or species-specific miRNAs are involved.

### Evaluation of miRNA targeting criteria in non-model organisms

Current miRNA targeting rules, inferred on interactions, mostly consisting of conserved miRNAs from *A. thaliana* [9], have been applied to other species for target prediction [28–31]. However, to the best of our knowledge, no comprehensive investigation into the suitability of these fixed targeting criteria has been performed in non-model organisms. The characterization of miRNA-mRNA interactions has been facilitated by both the increased complexity of experiments involving non-model plant species and through the analysis of RNA degradation profiles (PARE [3] sequencing and more recently NanoPARE [32]), which despite technical limitations (e.g. sequencing bias) can provide reliable high-throughput pseudo-validation of microRNA-mediated cleavage sites.

To investigate the suitability and portability of the fixed *Allen et al*. rules on non-model organisms and evaluate the scope for customised, organism-specific rules, we conducted an exploratory analysis using as input the HC degradome-supported miRNA-mRNA interactions reported by PAREameters. We compared the inferred rules for flower and leaf tissues in several organisms to produce a quantitative summary of the variation ranges of thresholds for the selected rules. Table 4 shows these summaries of inferred criteria per organism; Figure 3A illustrates the position-specific distributions of G:U pairs, mismatches and gaps, and Figure 3B shows the MFE ratio distributions for the miRNA-mRNA duplexes from flower tissue across organisms in *A. thaliana*, *A. trichopoda*, *O. sativa* and *T. aestivum*. Similar plots for leaf tissue in *A. thaliana*, *A. trichopoda* and *G. max* are presented in SF2.

**Table 4.**
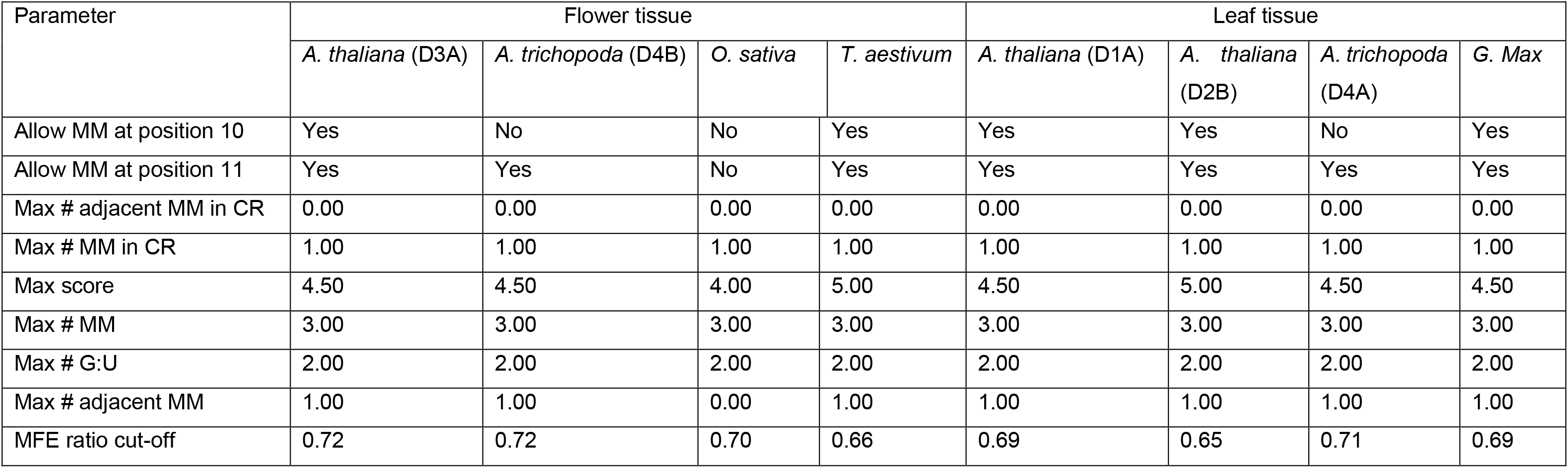
Overview of data-inferred thresholds inferred using PAREameters on model and non-model organisms in flower and leaf tissue. Differences are observed both between organisms (e.g. monocots vs dicots) and between tissues.

| Parameter | Flower tissue |  |  |  | Leaf tissue |  |  |  |
| --- | --- | --- | --- | --- | --- | --- | --- | --- |
|  | *A. thaliana* (D3A) | *A. trichopoda* (D4B) | *O. sativa* | *T. aestivum* | *A. thaliana* (D1A) | *A. thaliana* (D2B) | *A. trichopoda* (D4A) | *G. Max* |
| Allow MM at position 10 | Yes | No | No | Yes | Yes | Yes | No | Yes |
| Allow MM at position 11 | Yes | Yes | No | Yes | Yes | Yes | Yes | Yes |
| Max # adjacent MM in CR | 0.00 | 0.00 | 0.00 | 0.00 | 0.00 | 0.00 | 0.00 | 0.00 |
| Max # MM in CR | 1.00 | 1.00 | 1.00 | 1.00 | 1.00 | 1.00 | 1.00 | 1.00 |
| Max score | 4.50 | 4.50 | 4.00 | 5.00 | 4.50 | 5.00 | 4.50 | 4.50 |
| Max # MM | 3.00 | 3.00 | 3.00 | 3.00 | 3.00 | 3.00 | 3.00 | 3.00 |
| Max # G:U | 2.00 | 2.00 | 2.00 | 2.00 | 2.00 | 2.00 | 2.00 | 2.00 |
| Max # adjacent MM | 1.00 | 1.00 | 0.00 | 1.00 | 1.00 | 1.00 | 1.00 | 1.00 |
| MFE ratio cut-off | 0.72 | 0.72 | 0.70 | 0.66 | 0.69 | 0.65 | 0.71 | 0.69 |

**Figure 3:**
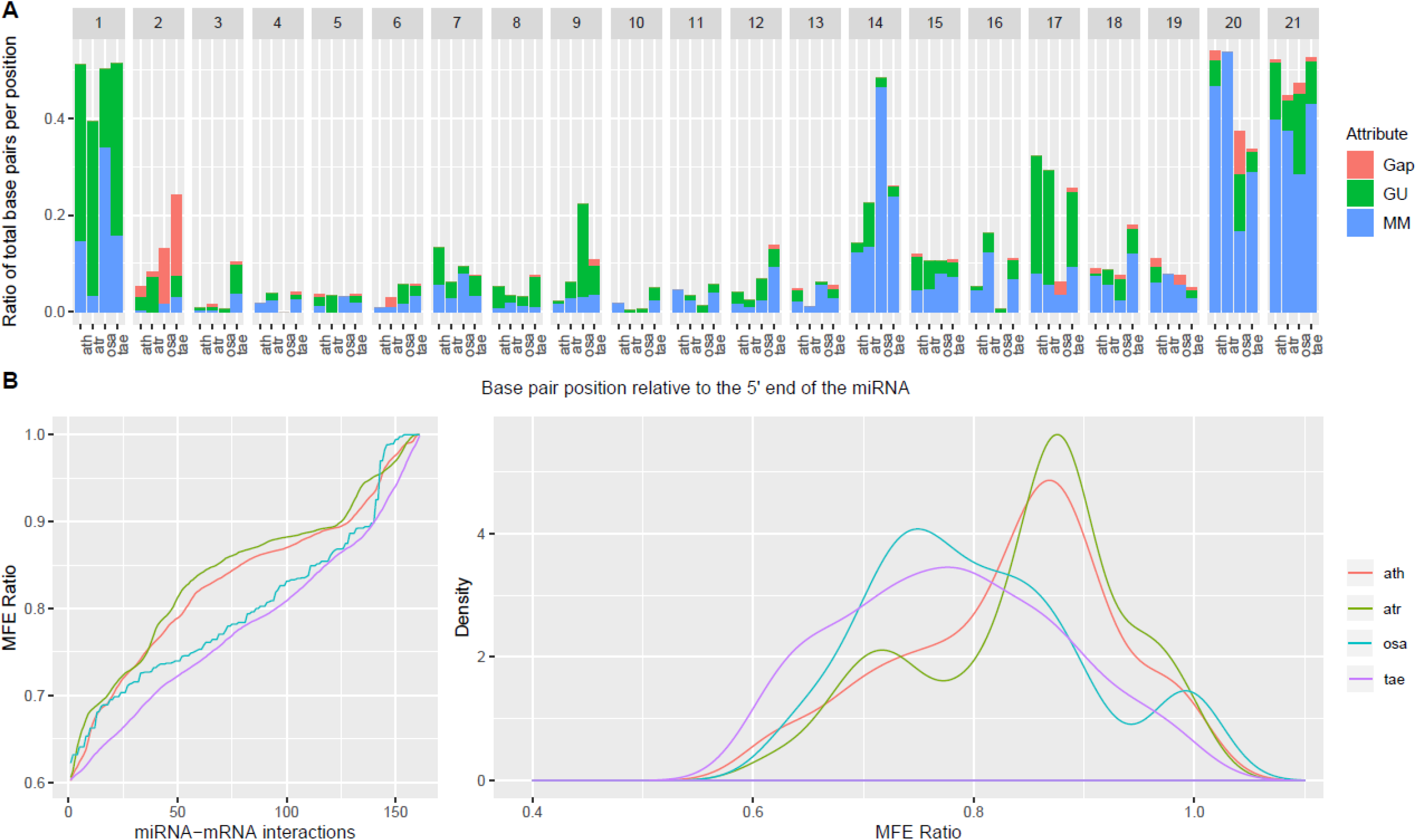
Side-by-side comparison of flower miRNA-mRNA interaction property distributions in monocots and dicots. The position-specific properties (panel A) and MFE ratio distribution (panel B) of miRNA-mRNA interactions from flower tissues in *A. thaliana*, *A. amborella*, *O. sativa* and *T. aestivum*. The position-specific properties showed significant differences at certain positions and there is a clear separation in the MFE distributions between monocots and dicots. The differences in properties for particular organisms from the current *A. thaliana* inferred criteria (e.g. MMs at position 1 for *A. amborella*, G:U pairs at position 9 for *O. sativa*, and almost perfect complementarity at positions 16-17 for *O. sativa*) and the distinction observed for the MFE ratios support the hypothesis that species or tissue specific, and data-inferred criteria may reveal a more accurate set of miRNA-mRNA regulatory interactions.

The distributions of position-specific properties in flower tissue show interesting variations between species. To evaluate whether the non-model organism distributions differ from the *A. thaliana* distributions, we used the offset χ^2^ test and a localized Fisher’s exact test (Table 5). The former show significant differences at position 1, positions 1, 2, 9, 14, 17 and 20, and positions 2, 3 and 20, for *A. trichopoda*, *O. sativa* and *T. aestivum*, respectively. The results of the localized Fisher’s exact test show significant differences between mismatches at position 1, positions 1, 14, and 20, and positions 14 and 20, for *A. trichopoda*, *O. sativa* and *T. aestivum*, respectively, supporting the conclusion that species specific, data driven criteria could facilitate a better description of the miRNA-mRNA. interactions. Additionally, the tests show significant differences between G:U pairs at positions 1, 7, 9 and 17 in *O. sativa* and significant differences in gaps at position 2 for both *O. sativa* and *T. aestivum*.

**Table 5:** χ^2^ and Fisher’s exact test significance results on the position-specific properties for non-model organisms versus *A. thaliana* in flower tissue. Values before the significance threshold (*p*-value ≤ 0.05) are highlighted in bold. The first position and the 9^th^ suggest that subtle differences do exist between *A. thaliana* and other organisms in key positions that determine the selection and mode of action for miRNAs.

| Species | *A. trichopoda* |  |  |  | *O. sativa* |  |  |  | *T. aestivum* |  |  |  |
| --- | --- | --- | --- | --- | --- | --- | --- | --- | --- | --- | --- | --- |
| Position | $\chi^2$ | MM | GU | Gap | $\chi^2$ | MM | GU | Gap | $\chi^2$ | MM | GU | Gap |
| 1 | **0.03** | **0.00** | 1.00 | 1.00 | **0.00** | **0.00** | **0.00** | 1.00 | 0.99 | 1.00 | 1.00 | 1.00 |
| 2 | 0.66 | 1.00 | 0.37 | 1.00 | **0.03** | 0.62 | 0.36 | **0.02** | **0.00** | 0.36 | 1.00 | **0.00** |
| 3 | 1.00 | 1.00 | 1.00 | 1.00 | 0.95 | 0.99 | 1.00 | 0.99 | **0.05** | 0.21 | 0.06 | 0.99 |
| 4 | 0.95 | 1.00 | 0.62 | 0.99 | 0.79 | 0.62 | 1.00 | 1.00 | 0.91 | 0.99 | 1.00 | 1.00 |
| 5 | 0.92 | 0.62 | 1.00 | 0.99 | 0.64 | 0.68 | 0.36 | 0.99 | 0.94 | 1.00 | 1.00 | 1.00 |
| 6 | 0.79 | 1.00 | 1.00 | 0.62 | 0.39 | 1.00 | 0.21 | 1.00 | 0.61 | 0.44 | 0.36 | 0.99 |
| 7 | 0.38 | 0.53 | 0.25 | 1.00 | 0.18 | 0.61 | **0.05** | 1.00 | 0.55 | 0.53 | 0.40 | 1.00 |
| 8 | 0.68 | 1.00 | 0.44 | 1.00 | 0.91 | 1.00 | 0.72 | 1.00 | 0.94 | 1.00 | 0.56 | 0.99 |
| 9 | 0.56 | 0.99 | 0.36 | 1.00 | **0.00** | 0.99 | **0.00** | 1.00 | 0.12 | 0.72 | 0.06 | 1.00 |
| 10 | 0.79 | 0.62 | 1.00 | 1.00 | 0.71 | 0.62 | 1.00 | 1.00 | 0.79 | 1.00 | 0.36 | 0.99 |
| 11 | 0.84 | 0.72 | 1.00 | 1.00 | 0.38 | 0.21 | 1.00 | 1.00 | 0.79 | 1.00 | 0.62 | 1.00 |
| 12 | 0.93 | 1.00 | 1.00 | 1.00 | 0.9 | 1.00 | 0.49 | 0.99 | 0.15 | 0.08 | 0.72 | 1.00 |
| 13 | 0.73 | 0.68 | 0.36 | 0.99 | 0.61 | 0.33 | 0.68 | 1.00 | 0.92 | 1.00 | 1.00 | 1.00 |
| 14 | 0.22 | 0.83 | 0.04 | 0.99 | **0.00** | **0.00** | 1.00 | 1.00 | 0.19 | **0.04** | 1.00 | 1.00 |
| 15 | 0.98 | 0.99 | 0.79 | 0.99 | 0.33 | 0.40 | 0.21 | 1.00 | 0.50 | 0.56 | 0.25 | 1.00 |
| 16 | 0.10 | 0.05 | 0.44 | 1.00 | 0.43 | 0.21 | 1.00 | 1.00 | 0.53 | 0.56 | 0.44 | 1.00 |
| 17 | 0.95 | 0.61 | 1.00 | 1.00 | **0.00** | 0.40 | **0.00** | 0.62 | 0.51 | .001 | 0.16 | 0.99 |
| 18 | 0.49 | 0.79 | 0.36 | 1.00 | 0.12 | 0.13 | 0.21 | 1.00 | 0.18 | 0.49 | 0.11 | 1.00 |
| 19 | 0.37 | 0.79 | 0.36 | 0.62 | 0.59 | 1.00 | 0.21 | 1.00 | 0.53 | 0.53 | 0.99 | 0.62 |
| 20 | 0.16 | 0.39 | 0.11 | 0.62 | **0.00** | **0.00** | 0.14 | 0.08 | **0.02** | **0.01** | 0.99 | 0.62 |
| 21 | 0.44 | 0.88 | 0.23 | 1.00 | 0.28 | 0.10 | 0.43 | 0.62 | 0.83 | 0.77 | 0.65 | 1.00 |

Next to the position-specific properties, the MFE ratio (part of the targeting criteria) was also investigated as a discriminative feature (Figure 3B); the Kolmogorov-Smirnov test was used to evaluate differences between distributions of different species. The distribution of MFE ratios and results of the statistical test, presented in Table 6, illustrate the differences between monocots and dicots, with significant differences only reported when comparing different groups.

**Table 6:** **Results of the Kolmogorov-Smirnov test when evaluating the differences between MFE ratio distributions** of HC miRNA-mRNA interactions found in flower tissue in model and non-model organisms. Values before the significance threshold (*p*-value ≤ 0.05) are highlighted in bold. The results highlight the significant differences observed between dicots (*A. thaliana* and *A. trichopoda*) and monocots (*O. sativa* and *T. aestivum*).

| Species | *A. thaliana* | *A. trichopoda* | *O. sativa* | *T. aestivum* |
| --- | --- | --- | --- | --- |
| *A. thaliana* | 1 | 0.331 | **2.54×10<sup>-4</sup>** | **6.89×10<sup>-6</sup>** |
| *A. trichopoda* |  | 1 | **1.93×10<sup>-7</sup>** | **1.38×10<sup>-8</sup>** |
| *O. sativa* |  |  | 1 | 0.166 |
| *T. aestivum* |  |  |  | 1 |

To evaluate the differences in number and identity of predicted miRNA targets when using the *Allen et al.* and PAREameters inferred criteria on the non-model organisms, we performed target prediction using PAREsnip2. The inferred criteria were able to capture a larger number of interactions; the only exception was observed for the D6 (*O. sativa*) dataset for which 149 interactions from 42 miRNAs were found using the *Allen et al.* rules and 115 interactions from 33 miRNAs using the inferred rules with an overlap of 100%. The larger number of interactions reported for the D5 (*G. max*) and D7 (*T. aestivum*) datasets when compared to D4 (*A. trichopoda*) and D6 (*O. sativa*), may have arisen from number of repeat regions or duplicated transcripts present within the current genome annotation.

Lastly, we investigated the overlap between the miRNAs and their interactions for each set of criteria, presented in Table 7, and concluded that, with the exception of D6 (O*. sativa*), a higher number of miRNAs and their interactions were specific to the inferred criteria, highlighting yet again the distance from the *Allen et al.* criteria. For this analysis we used the default retain rate of 0.85. To explore its effect on the overlap between the *Allen et al.* targeting criteria and the inferred criteria, we repeated the analysis using a retain rate value of 1, to capture all PAREameters reported HC interactions. All of the *Allen et al.* captured interactions were a subset of the interactions captured by the PAREameters inferred criteria when using a retain rate of 1 (ST12); the increase in miRNAs with targets varies between 4 (D6) and 102 (D7) and the increase in reported interactions varies between 12 (D6) and 783 (D7), depending on the organism/dataset in question. These results further suggest that the *Allen et al.* criteria may have been too stringent, or inadequately calibrated for the specific organism or miRNAs in question.

**Table 7:** Intersection analysis of interactions predicted using either the *Allen et al.* rules or the PAREameters inferred rules on various datasets. The number of interactions reported by PAREsnip2 using the *Allen et al.* criteria and the PAREameters inferred criteria on the non-model organisms varies between organisms and tissues; the number in brackets represents the miRNAs and interactions specific to the criteria used and could be used to approximate the accuracy of the prediction on non-model organisms. The exact sensitivity and precision values cannot be computed on non-model organisms due to the lack of a large enough set of validated interactions.

| Dataset | *Allen et al.* miRNAs | *Allen et al.* interactions | Inferred miRNAs | Inferred interactions |
| --- | --- | --- | --- | --- |
| D4A | 70 (3) | 203 (9) | 72 (5) | 210 (16) |
| D4B | 66 (2) | 174 (4) | 68(4) | 182 (12) |
| D5 | 143 (6) | 2118 (64) | 143 (6) | 2243 (189) |
| D6 | 42 (9) | 149 (34) | 33 (0) | 115 (0) |
| D7 | 91 (2) | 1257 (50) | 99 (10) | 1417 (210) |

## DISCUSSION

Larger, more diverse experiments, outside of the context of model organisms rely on computational methods, that extract features from sequencing data, for achieving a comparable sensitivity with well-studied organisms such as *A. thaliana*. The recent changes implemented for miRNA classification criteria [6–8] have signalled the need for a revised approach for defining their mode of action too.

In this paper, we describe PAREameters, a novel approach and pipeline that enables data-driven inference of plant miRNA targeting criteria, which ensures maximal usage of input data (degradome datasets and corresponding sRNA samples). Through refining the targeting criteria, the discovery and characterization of new miRNA-mRNA interactions per tissue or organism (both model and non-model) becomes possible. When evaluating the performance of the PAREameters inferred criteria, we observed an increase in sensitivity compared to the *Allen et al.* criteria [9] over all the *A. thaliana* datasets, whilst also maintaining precision on most datasets, when benchmarked against a set of experimentally validated miRNA-mRNA interactions.

The comparison of validated miRNA-mRNA interaction properties between conserved and species-specific miRNAs highlighted interesting results. When investigating the features of conserved miRNA interactions, we observed similar patterns to that of *Allen et al.* [9] regarding complementarity in the core region of the miRNA (2-13) and also at the canonical position 10. This observation is further supported by a recent study of highly conserved miRNAs in *N. benthamiana* [33], where it was shown that a single mismatch at the 5’ end of miR160 significantly diminished target site efficacy, and two or more consecutive mismatches at the 5′ end fully abolished it. Furthermore, the authors highlighted that a single nucleotide mismatch at positions 9 and 10, in addition to combinations of mismatches at positions 9, 10, and 11 led to the complete elimination of the responsiveness of miR164. However, the species-specific miRNAs tended to tolerate more flexibility at these positions.

Through our study, we highlight that targeting criteria inferred on subsets of interactions are less compatible with one another and often lead to a considerable drop in sensitivity. Given the current understanding of the miRNA-mRNA interactions in various species, it is difficult to propose a biological interpretation of these variations, however we can conclude that a customised selection of thresholds may result in a more detailed overview of regulatory interactions and facilitate a more in-depth assessment of the underlying regulatory networks. Furthermore, the differences observed in the flower tissue between monocots and dicots emphasise the usefulness of data-inferred, species/tissue specific thresholds.

The tool is optimized both in runtime and computational resource usage; the analysis of a typical *A. thaliana* and *T. aestivum* sample completes in ~30 minutes and 1 day 10 hours, with 6GB and 10GB memory (RAM) requirements, respectively. PAREameters was also implemented as a user-friendly, cross-platform (Windows, Linux and MacOS) application that enables users to analyse sequencing datasets without the need of specialized support or dedicated hardware. These features recommend it for a wide variety of experimental designs and organisms that will enable further understanding of the subtle variations in miRNA-mRNA interactions in different species, tissues and treatments. In addition, the novel data-driven approach may enable new discoveries within the RNA silencing pathways.

## AVAILABILITY

PAREameters is available as part of the UEA sRNA Workbench [16]; it can be downloaded from http://srna-workbench.cmp.uea.ac.uk/. The source code has been released on GitHub and is accessible at https://github.com/sRNAworkbenchuea/UEA_sRNA_Workbench/.

## Supporting information

Tutorial data

