## Supplemental Figures for "PAREameters: computational inference of plant microRNA-mRNA targeting rules using RNA sequencing data"

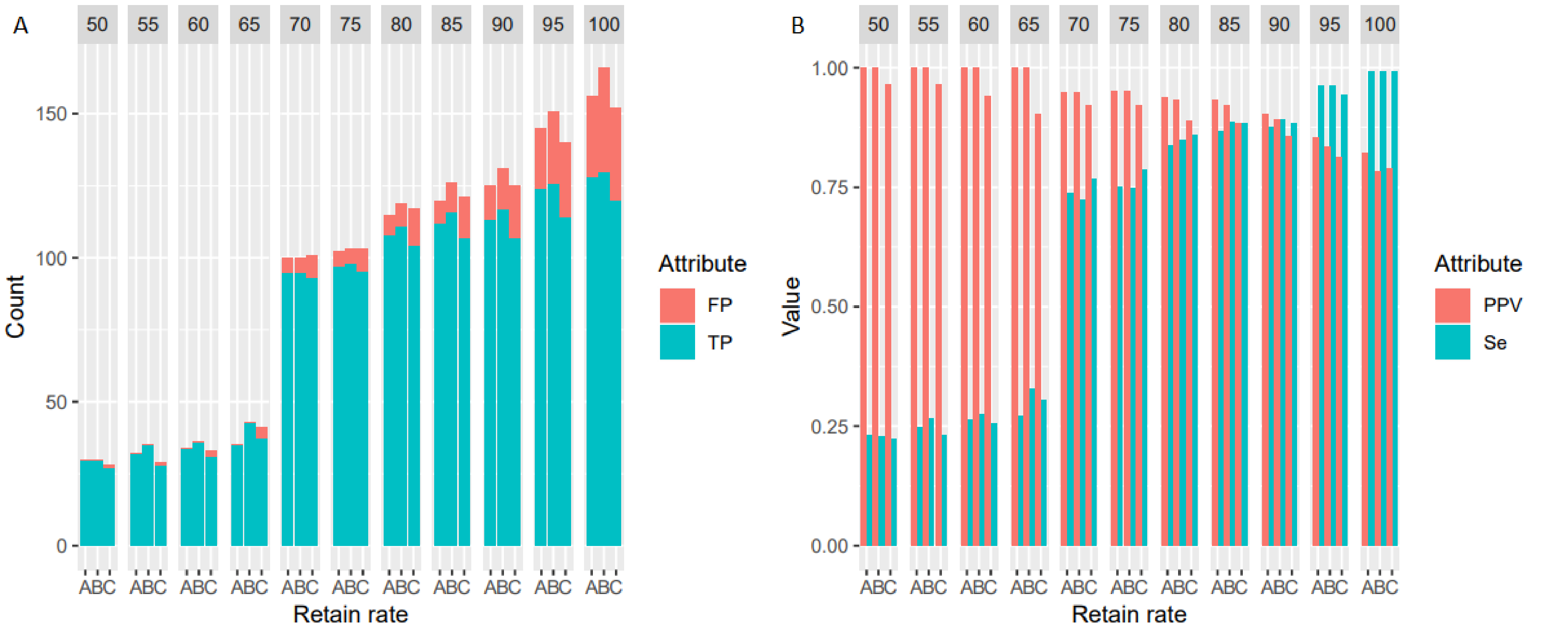


Supplementary figure 1: **Accuracy (Sensitivity/PPV) dependence on the retain rate parameter**. Clustered frequency histograms of predicted interactions vs validated/non-validated ones (panel A) and the variation in sensitivity and precision values for increasing values of the retain rate parameter on three A. *Thaliana* leaf replicates, the D1 dataset (panel B) highlight the existence of a data-driven optimum for the retain rate parameter. For this particular dataset the optimum on the Se/PPV ratio is achieved for 0.85. The data-driven optimal value for this parameter is suggested based on the input; however, it still remains a user-configurable parameter.

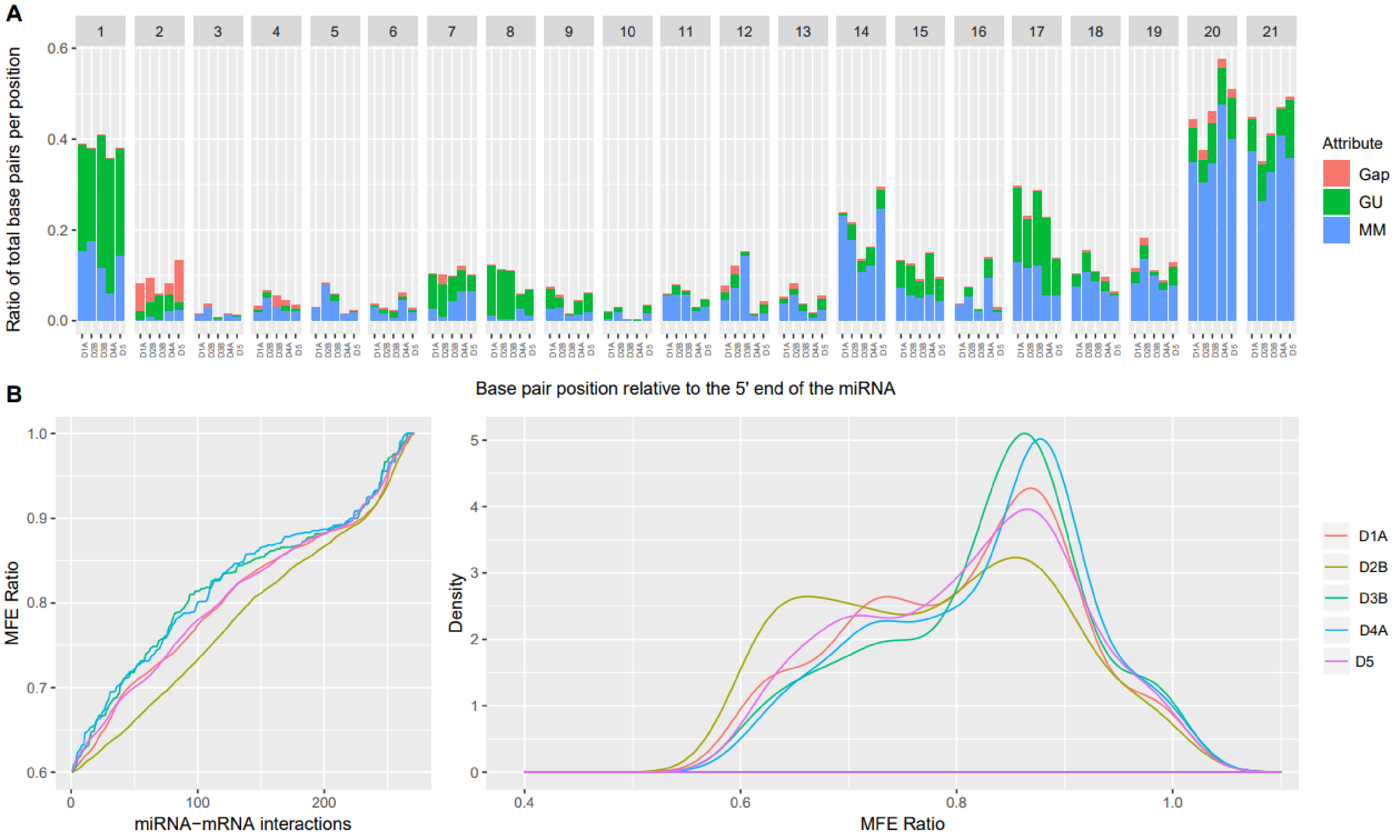


Supplementary figure 2: **Side-by-side comparison of leaf miRNA-mRNA interaction property distributions in different species and datasets.** The position-specific properties (panel A) and MFE ratio distribution (panel B) of miRNA-mRNA interactions from leaf tissues in *A. thaliana*, *A. amborella* and *G. max*. The differences in properties for particular organisms and the differences observed for the MFE ratios support the hypothesis that species or tissue specific, and data-driven criteria may reveal a more accurate set of miRNA-mRNA regulatory interactions.
