## Supplemental Tables 1-8 for "PAREameters: computational inference of plant microRNA-mRNA targeting rules using RNA sequencing data"

| Parameter | *Allen et al.* | Manually inferred | Permissive PAREameters criteria |
|---|---|---|---|
| Allow MM at position 10 | No | Yes | Yes |
| Allow MM at position 11 | No | Yes | Yes |
| Max # adjacent MM in CR | 1 | 0 | 2 |
| Max # MM in CR | 2 | 2 | 3 |
| Max score | 4 | 5 | 6 |
| Max # MM | 4 | 4 | 6 |
| Max # G:U | 4 | 3 | 5 |
| Max # adjacent MM | 2 | 1 | 4 |
| MFE ratio cut-off | 0.7 | 0.65 | 0.6 |

Supplementary table 1: **The parameter values for the *Allen et al.*, manually inferred and PAREameters permissive criteria, derived from the experimentally validated *A. thaliana* miRNA-mRNA interactions.** The *Allen et al.* criteria were inferred in 2005 on a small set of experimentally validated interactions, in *A. Thaliana*; the manually inferred criteria were inferred on a larger set of experimentally validated interactions, for the same organism. The permissive parameters are used initially by PAREameters to find HC interactions. The inferred criteria are then extracted from those using the retain rate parameter.

| Dataset | # miRNAs | # Validated in dataset | Allen V | Inferred V | Allen NV | Inferred NV | Allen Se | Inferred Se | Allen PPV | Inferred PPV | Se gain | PPV difference |
|---|---|---|---|---|---|---|---|---|---|---|---|---|
| D1A | 37 | 129 | 105 | 122 | 9 | 20 | 81.40% | 94.57% | 92.11% | 85.92% | 13.18% | -6.19% |
| D1B | 38 | 131 | 109 | 126 | 11 | 25 | 83.21% | 96.18% | 90.83% | 83.44% | 12.98% | -7.39% |
| D1C | 35 | 121 | 95 | 115 | 12 | 25 | 78.51% | 95.04% | 88.79% | 82.14% | 16.53% | -6.64% |
| D2A | 40 | 140 | 117 | 133 | 14 | 43 | 83.57% | 95.00% | 89.31% | 75.57% | 11.43% | -13.75% |
| D2B | 38 | 137 | 113 | 129 | 13 | 43 | 82.48% | 94.16% | 89.68% | 75.00% | 11.68% | -14.68% |
| D2C | 40 | 144 | 117 | 138 | 3 | 11 | 81.25% | 95.83% | 97.50% | 92.62% | 14.58% | -4.88% |
| D3A | 32 | 79 | 64 | 77 | 4 | 10 | 81.01% | 97.47% | 94.12% | 88.51% | 16.46% | -5.61% |
| D3B | 29 | 70 | 57 | 66 | 11 | 17 | 81.43% | 94.29% | 83.82% | 79.520% | 12.86% | -4.31% |
| D3C | 36 | 111 | 84 | 106 | 6 | 12 | 75.68% | 95.50% | 93.33% | 89.83% | 19.82% | -3.50% |
| D3D | 35 | 104 | 88 | 101 | 3 | 7 | 84.62% | 97.12% | 96.70% | 93.52% | 12.50% | -3.19% |

Supplementary table 2: **Sensitivity and precision values for the *Allen et al.* and manually inferred criteria over all the *A. Thaliana* datasets.** Allen = *Allen et al.* rules, V = validated, NV = non-validated, Se = sensitivity and PPV = precision. An increase in the achieved Se is observed for the inferred criteria; the decrease in PPV is mainly due to the lack of low-throughput validations for a subset of interactions, for. which a clear signal is observed in the PARE datasets.

| Transition | D1A | | | D1B | | | D1C | | |
|---|---|---|---|---|---|---|---|---|---|
| | Sensitivity gain | Precision loss | Ratio | Sensitivity gain | Precision loss | Ratio | Sensitivity gain | Precision loss | Ratio |
| 0.50-0.55 | 1.55% | 0.00% | - | 3.82% | 0.00% | - | 0.83% | 0.12% | 6.710 |
| 0.55-0.60 | 1.55% | 0.00% | - | 0.76% | 0.00% | - | 2.48% | -2.61% | 0.95 |
| 0.60-0.65 | 0.78% | 0.00% | - | 5.34% | 0.00% | - | 4.96% | -3.70% | 1.34 |
| 0.65-0.70 | 46.51% | -5.00% | 9.30 | 39.69% | -5.00% | 7.94 | 46.28% | 1.84% | 25.22 |
| 0.70-0.75 | 1.55% | 0.10% | 15.81 | 2.29% | 0.15% | 15.73 | 1.65% | 0.15% | 10.75 |
| 0.75-0.80 | 8.53% | -1.18% | 7.20 | 9.92% | -1.87% | 5.31 | 7.44% | -3.34% | 2.22 |
| 0.80-0.85 | 3.10% | -0.58% | 5.35 | 3.82% | -1.21% | 3.14 | 2.48% | -0.46% | 5.40 |
| 0.85-0.90 | 0.78% | -2.93% | 0.26 | 0.76% | -2.75% | 0.28 | 0.00% | -2.83% | 0.00 |
| 0.90-0.95 | 8.53% | -4.88% | 1.75 | 6.87% | -5.87% | 1.17 | 5.79% | -4.17% | 1.37 |
| 0.95-1 | 3.10% | -3.47% | 0.89 | 3.05% | -5.13% | 0.60 | 4.96% | -2.48% | 2.00 |

Supplementary table 3: **Sensitivity gain vs precision loss and the absolute ratio between them for values of the retain rate parameter in the 0.50-1.00 range, over three *A. thaliana* leaf replicates (D1 dataset).** The sensitivity and precision were evaluated on the experimentally validated miRNA-mRNA interactions in *A. thaliana* that had corresponding HC transcript peaks within the degradome dataset. The ratio is defined as the absolute value of sensitivity/precision. The optimal value is obtained for the last transition that maximises sensitivity gain whilst simultaneously minimising precision loss i.e. the stopping criteria is achieved for the minimum value of the absolute ratio. For this dataset, the value is achieved for the 0.85-0.90 transition, indicating that the retain rate parameter should be set at 0.85.

| Dataset | Raw reads | Unique reads | Genome matched | % matched | miRNAs present |
|---|---|---|---|---|---|
| D1A | 6664998 | 1342846 | 1006022 | 74.92% | 230 |
| D1B | 4492236 | 1093344 | 818760 | 74.89% | 213 |
| D1C | 5148552 | 1106222 | 837919 | 75.75% | 230 |
| D2A | 27870710 | 1935025 | 1091177 | 56.40% | 252 |
| D2B | 26211889 | 908368 | 525503 | 57.90% | 239 |
| D2C | 28700595 | 568633 | 240613 | 42.31% | 209 |
| D3A | 18460973 | 3797561 | 2534869 | 66.75% | 200 |
| D3B | 10408796 | 1633730 | 1101861 | 67.44% | 186 |
| D3C | 19946757 | 2837304 | 1647286 | 58.06% | 211 |
| D3D | 6645127 | 1176424 | 836656 | 71.12% | 141 |
| D4A | 18092450 | 1765893 | 656968 | 37.20% | 153 |
| D4B | 25781233 | 4560684 | 2685317 | 58.88% | 161 |
| D5 | 33230948 | 2370300 | 1641914 | 69.27% | 309 |
| D6 | 4029462 | 1991942 | 1991720 | 99.99% | 227 |
| D7 | 12541386 | 5178587 | 4325043 | 83.51% | 168 |

Supplementary table 4A: **Summary statistics of the sRNA sequencing data.** The number of raw (redundant), unique and genome matched reads. In order for an annotated miRNA to be considered present, it must have had an abundance >=5 within the library.

| Dataset | Raw reads | Unique reads | Transcriptome matched (+ strand) | % matched |
|---|---|---|---|---|
| D1A | 44871978 | 11114549 | 8802080 | 79.19% |
| D1B | 34315808 | 10103690 | 8176679 | 80.93.% |
| D1C | 25588818 | 7715251 | 6251721 | 81.03% |
| D2A | 115224802 | 19930692 | 10565361 | 53.01% |
| D2B | 107999423 | 19470487 | 11187427 | 57.46% |
| D2C | 191294550 | 7275123 | 1734835 | 23.85% |
| D3A | 5263291 | 2463251 | 2131493 | 86.53% |
| D3B | 4809175 | 2300541 | 1998764 | 86.88% |
| D3C | 12666325 | 3975280 | 3454112 | 86.89% |
| D3D | 53840936 | 9032093 | 6824080 | 75.56% |
| D4A | 19990216 | 4305009 | 1940976 | 45.09% |
| D4B | 12609502 | 3992618 | 1679932 | 42.08% |
| D5 | 26251057 | 7704474 | 6230026 | 80.86% |
| D6 | 4426044 | 2505523 | 2268297 | 90.53% |
| D7 | 35477509 | 14363576 | 10366761 | 72.17% |

Supplementary table 4B: **Summary statistics of the degradome sequencing data.** The number of raw (redundant), unique and transcriptome (positive strand only) matching reads within each degradome library.

| Parameter | D1A | D1B | D1C | D2A | D2B | D2C | D3A | D3B | D3C | D3D |
|---|---|---|---|---|---|---|---|---|---|---|
| Allow MM at position 10 | Yes | Yes | Yes | Yes | Yes | Yes | Yes | Yes | Yes | Yes |
| Allow MM at position 11 | Yes | Yes | Yes | Yes | Yes | Yes | Yes | Yes | Yes | Yes |
| Max number of adjacent MM in CR | 0 | 0 | 0 | 0 | 0 | 0 | 0 | 0 | 0 | 0 |
| Max number of MM in CR | 1 | 1 | 1 | 1 | 1 | 1 | 1 | 1 | 1 | 1 |
| Max score | 4.50 | 4.50 | 5.00 | 5.00 | 5.00 | 4.50 | 4.00 | 4.50 | 4.50 | 4.50 |
| Max number of MM | 3 | 3 | 3 | 3 | 3 | 3 | 3 | 3 | 3 | 3 |
| Max number of G:U | 2 | 2 | 2 | 2 | 2 | 2 | 2 | 2 | 2 | 2 |
| Max number of adjacent MM | 1 | 1 | 1 | 1 | 1 | 1 | 1 | 1 | 1 | 1 |
| MFE ratio cut-off | 0.69 | 0.70 | 0.69 | 0.66 | 0.65 | 0.67 | 0.72 | 0.71 | 0.70 | 0.71 |

Supplementary table 5: **The PAREameters inferred criteria for each of the Arabidopsis datasets.** MM = mismatch, CR = core region (2-13nt of miRNA) and MFE = minimum free energy.

| Dataset | Unique sRNAs | Unique PARE seqs | Run Time (hh:mm:ss) | Memory (GB) |
|---|---|---|---|---|
| D1A | 1342846 | 11114549 | 00:18:51 | 6 |
| D1B | 1093344 | 10103690 | 00:17:41 | 6 |
| D1C | 1106222 | 7715251 | 00:17:11 | 6 |
| D2A | 1935025 | 19930692 | 00:31:18 | 8 |
| D2B | 908368 | 19470487 | 00:38:09 | 8 |
| D2C | 568633 | 7275123 | 01:04:24 | 8 |
| D3A | 3797561 | 2463251 | 00:18:15 | 5 |
| D3B | 1633730 | 2300541 | 00:16:52 | 5 |
| D3C | 2837304 | 3975280 | 00:20:30 | 5 |
| D3D | 1176424 | 9032093 | 00:17:59 | 5 |
| D4A | 1765893 | 4305009 | 01:30:15 | 6 |
| D4B | 4560684 | 3992618 | 01:34:12 | 6 |
| D5 | 2370300 | 7704474 | 02:00:26 | 7 |
| D6 | 1991942 | 2505523 | 00:41:43 | 5 |
| D7 | 5178587 | 14363576 | 34:37:10 | 10 |

Supplementary table 6: **The timing and memory usage results for the PAREameters analysis of the Arabidopsis datasets.** The size of the input (as unique entries) and complexity of the underlying genome are the main drivers for both run-time and resource usage.

| Dataset | Test Inferred SE | Test Inferred PPV |
|---|---|---|
| D1A | 0.81 | 0.83 |
| D1B | 0.78 | 0.83 |
| D1C | 0.80 | 0.82 |
| D2A | 0.79 | 0.78 |
| D2B | 0.77 | 0.87 |
| D2C | 0.67 | 0.89 |
| D3A | 0.68 | 0.78 |
| D3B | 0.77 | 0.75 |
| D3C | 0.78 | 0.96 |
| D3D | 0.75 | 1.000 |

Supplementary table 7: **The median sensitivity and PPV (precision) values for the cross-validation experiments on the Arabidopsis datasets.** The CV was done on a 75/25% split for training and testing, respectively. Each analysis was repeated 50 times and the median value was recorded.

| Training size (% of total) | D1A Sensitivity | D1A Precision | D1B Sensitivity | D1B Precision | D1C Sensitivity | D1C Precision |
|---|---|---|---|---|---|---|
| 0.1 | 68.97% | 95.17% | 48.72% | 97.22% | 71.76% | 90.91% |
| 0.2 | 70.87% | 93.80% | 59.62% | 94.77% | 66.67% | 90.42% |
| 0.3 | 77.22% | 92.41% | 64.29% | 94.46% | 73.81% | 89.42% |
| 0.4 | 79.22% | 91.54% | 75.64% | 91.91% | 70.83% | 88.15% |
| 0.5 | 78.13% | 89.92% | 76.92% | 89.71% | 77.50% | 86.44% |
| 0.6 | 75.49% | 87.95% | 77.88% | 87.50% | 77.08% | 84.09% |
| 0.7 | 78.95% | 85.29% | 76.92% | 85.33% | 77.78% | 83.83% |
| 0.8 | 82.00% | 80.77% | 80.77% | 83.33% | 79.17% | 82.35% |
| 0.9 | 83.33% | 78.17% | 84.62% | 79.29% | 79.17% | 80.00% |

Supplementary table 8: **The median sensitivity and precision values for the training-size experiment on the Arabidopsis datasets.** For each dataset, an increase in training-size resulting in an overall increase in sensitivity, supporting the conclusion that deeper sequencing experiments can yield more information on regulatory interactions. The characteristics of the sampling data, such as sequencing depth and complexity also influence the Se and PPV values (as observed for replicate B).

| Retain rate | D1A Sensitivity | D1A Precision | D1B Sensitivity | D1B Precision | D1C Sensitivity | D1C Precision |
|---|---|---|---|---|---|---|
| 0.5 | 23.26% | 100.00% | 22.90% | 100.00% | 22.31% | 96.43% |
| 0.55 | 24.81% | 100.00% | 26.72% | 100.00% | 23.14% | 96.55% |
| 0.6 | 26.36% | 100.00% | 27.48% | 100.00% | 25.62% | 93.94% |
| 0.65 | 27.13% | 100.00% | 32.82% | 100.00% | 30.58% | 90.24% |
| 0.7 | 73.64% | 95.00% | 72.52% | 95.00% | 76.86% | 92.08% |
| 0.75 | 75.19% | 95.10% | 74.81% | 95.15% | 78.51% | 92.23% |
| 0.8 | 83.72% | 93.91% | 84.73% | 93.28% | 85.95% | 88.89% |
| 0.85 | 86.82% | 93.33% | 88.55% | 92.06% | 88.43% | 88.43% |
| 0.9 | 87.60% | 90.40% | 89.31% | 89.31% | 88.43% | 85.60% |
| 0.95 | 96.12% | 85.52% | 96.18% | 83.44% | 94.21% | 81.43% |
| 1 | 99.22% | 82.05% | 99.24% | 78.31% | 99.17% | 78.95% |

Supplementary table 9: **Analysis of the retain rate parameter using the *A. Thaliana* D1 dataset.** The dual optimization problem for maximizing the Se and minimizing the PPV loss is solved using the Se/PPV ratio, which for this dataset achieves its minimum for 0.85.

| Dataset | *Allen et al.* miRNAs | *Allen et al.* interactions | Inferred miRNAs | Inferred interactions | Difference in miRNAs | Difference in interactions |
|---|---|---|---|---|---|---|
| D4A | 70 | 203 | 87 | 272 | 17 | 69 |
| D4B | 66 | 174 | 79 | 208 | 13 | 34 |
| D5 | 143 | 2118 | 190 | 2842 | 47 | 724 |
| D6 | 42 | 149 | 46 | 161 | 4 | 12 |
| D7 | 91 | 1257 | 193 | 2040 | 102 | 783 |

Supplementary table 12: **Frequency summary on the count of miRNAs and corresponding interactions determined using either the Allen et al rules or the inferred rules for non-model organisms.** The difference in number of miRNAs and their interactions when comparing the *Allen et al.* criteria and the criteria inferred by PAREameters using a retain rate of 1. All of the *Allen et al.* reported interactions are a subset of the inferred criteria reported interactions when using a retain rate of 1.
