## Supplementary material for "PAREameters: computational inference of plant microRNA-mRNA targeting rules using RNA sequencing data": Tutorial data: read_me.txt

This is the tutorial data for PAREameters. 

To perform an analysis using PAREameters, please download the latest version of the sRNA Workbench.

https://sourceforge.net/projects/srnaworkbench/files/latest/download

You can run PAREameters through the command line. 

An example of how to run PAREameters using the command-line and the tutorial data can be found below:

java -jar -Xmx8g Workbench.jar -tool pareameters -pare_file PAREameters_tutorial_data/degradome.fa -srna_file PAREameters_tutorial_data/miRNAs.fa -genome_file PAREameters_tutorial_data/TAIR10_chr1.fa -transcript_file PAREameters_tutorial_data/transcripts.fa -species_name Ath -output_dir pareameters_tutorial
