## Supplementary material for "PAREameters: computational inference of plant microRNA-mRNA targeting rules using RNA sequencing data": Tutorial data: user_guide.pdf

22/07/2019

User Guide

Joshua Thody, Vincent Moulton and Irina Mohorianu


#### Table of Contents


*Introduction* ... 2

*System requirements* ... 3

*Input data and parameters* ... 4

Input data ... 4

Tool configuration ... 4

Default parameters ... 5

*Launching PAREameters from the command-line (CMD)* ... 6

*Output* ... 6

Inferred targeting criteria and configurations ... 6

Predicted miRNA sequences with targets ... 7

High-confidence miRNA-mRNA interactions ... 7

Specific miRNA-mRNA interaction property files ... 7

Position specific properties ... 7
MFE ratio distribution ... 7
Base-pair properties ... 7
Cumulative alignment score distribution ... 8
Cumulative number of mismatches in the core region (2-13) ... 8
Cumulative number of G:U pairs in the core region (2-13) ... 8
Cumulative number of adjacent mismatches in the core region (2-13) ... 8
Cumulative number of mismatches ... 8
Cumulative number of G:U pairs ... 8
Cumulative number of adjacent mismatches ... 8

*Using R to produce plots* ... 8

Clustered histogram of position specific properties and Minimum free energy ratio distribution 9

### Introduction

MicroRNAs (miRNAs) are short, non-coding RNAs that influence the translation-rate of mRNAs by directing the RNA-induced silencing complex to sequence-specific targets. In plants, this typically results in cleavage and subsequent degradation of the target mRNA. This can be captured on a high-throughput scale using degradome sequencing, which supports miRNA target prediction by aligning degradation fragments to reference mRNAs enabling the identification of causal miRNA(s). The current criteria used for target prediction were inferred on experimentally validated *A. thaliana* interactions, which over-fitted a specific subset of miRNA interactions. In addition, the miRNA pathway in other organisms may have acquired specific changes, e.g. lineage specific miRNAs or new miRNA-mRNA interactions, and thus the previous criteria may not be optimal. PAREameters is a software tool for inferring plant miRNA targeting criteria from sequencing datasets, which can then be incorporated into sRNA target prediction analyses.

#### System requirements

The UEA sRNA Workbench has been tested on various platforms including:

- Mac OSX (Version 10.5 Leopard; 10.6 Snow Leopard; 10.7 Lion, 10.8 Mountain Lion, 10.9 Mavericks, 10.10 Yosemite, 10.13 High Sierra)
- Linux (Ubuntu Version 16.04)
- Windows 7 and 10

Currently, the software is built and tested on the official Oracle builds of Java only. However, most of the software should behave in the same way under open builds but we cannot guarantee this.

Required:

Java 8 (At the time of writing, later versions of Java are not yet supported)

Recommended:

Intel i5 quad core (or similar) 16GB RAM

### Input data and parameters

This section discusses the types of data that can be used to infer targeting criteria using the PAREameters pipeline. Additionally, we describe each parameter in detail.

#### Input data

- A small RNA and corresponding degradome dataset
- A reference genome from the organism in question
- A set of transcript sequences from the organism in question
- User defined miRNA sequences of interest (optional)

The sRNA and degradome libraries must be in redundant FASTA format with the adapters trimmed. FASTQ to FASTA and adapter removal tools are provided within the UEA sRNA Workbench. Additionally, any sequences containing ambiguous bases will be discarded as they cannot be accurately aligned.

#### Tool configuration

Below we describe the configurable parameters for the PAREamaters pipeline.

| Parameter | Possible values | Description |
|---|---|---|
| use_weighted_abundance | true/false | Use weighted fragment abundance (abundance/ # alignments) |
| min_sRNA_abundance | Integer >= 1 | The minimum sRNA abundance |
| min_tag_abundance | Integer >= 1 | The minimum tag abundance |
| min_sRNA_length | Integer >= 1 | The minimum sRNA length |
| max_sRNA_length | Integer <= 30 | The maximum sRNA length |
| min_tag_length | Integer >= 1 | The minimum tag length |
| max_tag_length | Integer <= 30 | The maxiumum tag length |
| retain_rate | double, [0,1] | The value for the retain rate parameter e.g. 0.75 |
| mfe_ratio_cutoff | double, [0,1] | The value for the MFE ratio cut-off e.g. 0.6 |
| category_0 | true/false | Allow category 0 peaks |
| category_1 | true/false | Allow category 1 peaks |
| category_2 | true/false | Allow category 2 peaks |
| category_3 | true/false | Allow category 3 peaks |
| category_4 | true/false | Allow category 4 peaks |
| filter_low_complexity_seqs | true/false | Filter low complexity sequences |

#### Default parameters

Below are the default parameters for the PAREameters pipeline.

| Parameter | Default value |
|---|---|
| use_weighted_abundance | false |
| min_sRNA_abundance | 5 |
| min_tag_abundance | 5 |
| min_sRNA_length | 19 |
| max_sRNA_length | 24 |
| min_tag_length | 19 |
| max_tag_length | 21 |
| retain_rate | 0.85 |
| mfe_ratio_cutoff | 0.65 |
| category_0 | true |
| category_1 | true |
| category_2 | false |
| category_3 | false |
| category_4 | false |
| filter_low_complexity_seqs | true |

### Launching PAREameters from the command-line (CMD)

In order to execute the sRNA Workbench and PAREsnip2 from the command line, navigate to the directory that you extracted the sRNA Workbench files to. Then type:

```
java -jar Workbench.jar -tool PAREameters
```

If no options are entered, the usage instructions will be printed to the command line. An example of a complete instruction is given below:

```
Usage:
  java [-XmxNg] -jar /path/to/Workbench.jar -tool pareameters -
pare_file path/to/pare/file -srna_file path/to/srna/file -
genome_file path/to/genome/file -transcript_file
path/to/transcript/file -output_dir path/to/output/directory –
species_name species_name [-mirna_file path/to/mirna/file] [-config
path/to/PAREameters/config/file] [-paresnip2_rules
path/to/paresnip2_rules/file]
```

For example, using the provided tutorial data, you would provide the following command:

```
java -jar Workbench.jar -tool pareameters -pare_file PAREameters_tutorial_data/degradome.fa
-srna_file PAREameters_tutorial_data/miRNAs.fa -genome_file PAREameters_tutorial_data/TAIR10_c
hr1.fa -transcript_file PAREameters_tutorial_data/transcripts.fa -species_name Ath -output_dir
 pareameters_test -mirna_file PAREameters_tutorial_data/ath_miRNAs.fa
```

Note: parameters in square brackets are optional. Default parameter files that can be used or edited are found in the default parameters directory of the Workbench. The species_name parameter **cannot** contain spaces; an example would be Arabidopsis. For any further analysis for the same species, please use the same species name, as this will speed up the analysis.

### Output

The output of the PAREameters file consists of the following:

1. A set of targeting criteria and configurations to be used for a sRNA target analysis
2. A set of predicted miRNA sequences with targets
3. A set of high-confidence miRNA-mRNA interactions
4. A collection of files specifying the properties of these miRNA-mRNA interactions

Below explain the provided output files in more detail.

#### Inferred targeting criteria and configurations

The PAREamaters pipeline will produce two files that can be used to help the user decide what parameters to use for their sRNA target prediction. In the provided form, these files can be used as input for a PAREsnip2 analysis. However, the provided parameters can also be used by another tool or custom script, if the configuration of parameters is permitted.

The “targeting_rules.txt” file contains the parameter values for each targeting criteria and the “parameters.txt” file contains the inferred MFE ratio.

#### Predicted miRNA sequences with targets

As part of the PAREameters pipeline, miRNAs are predicted using miRCat2 and miRPlant. These sequences are then provided to PAREsnip2, along with the corresponding degradome data, for target prediction. The “predicted_miRNAs_with_targets.fa” file contains the miRNA sequences, both known and new, that were predicted to also have targets within the degradome. This file is in non-redundant FASTA format.

#### High-confidence miRNA-mRNA interactions

The file “miRNA_alignments.csv” contains the PAREsnip2 results for those sequences that are predicted to be miRNAs. For a detailed description of the PAREsnip2 results, please see the PAREsnip2 user manual.

#### Specific miRNA-mRNA interaction property files

After completion of a PAREameters analysis, you will find a direction named “plot_csv_files” in the PAREameters output directory. This file contains all the recorded properties of the predicted miRNA-mRNA interactions, which are then used to infer targeting parameters. Using this information, it is possible to manually infer your own criteria that suits your required sensitivity and specificity.

##### Position specific properties

The “position_property_plot.csv” file contains the proportion of mismatches, G:U pairs and gaps at each position relative to the miRNA sequence. An R-script is provided to produce clustered histograms from this data, described later in this document.

##### MFE ratio distribution

The “mfe_ratio_plot.csv” file contains the cumulative distribution of the MFE ratios of the miRNA-mRNA interactions. An R-script is provided to produce the distribution plot and the density plot from this data, described later in this document.

##### Base-pair properties

The “basepair_property_plot.csv” file contains the proportion of possible base-pairs at each position (A:U, G:C, G:U, A:m, G:m, C:m, U:m, A:-, G:-, C:- and U:-).

##### Cumulative alignment score distribution

The “alignment_score_plot.csv” contains the cumulate distribution of alignment scores for the miRNA-mRNA interactions. This file can be used to decide an appropriate cut-off score for miRNA target prediction.

##### Cumulative number of mismatches in the core region (2-13)

The “core_mismatches_plot.csv” file contains the cumulative number of mismatches within the core region of all reported miRNA-mRNA interaction pairs.

##### Cumulative number of G:U pairs in the core region (2-13)

The “core_GU_pairs_plot.csv” file contains the cumulative number of G:U within the core region of all reported miRNA-mRNA interaction pairs.

##### Cumulative number of adjacent mismatches in the core region (2-13)

The “core_adj_mismatches_plot.csv” file contains the cumulative number of adjacent mismatches within the core region of all reported miRNA-mRNA interaction pairs.

##### Cumulative number of mismatches

The “mismatch_plot.csv” file contains the cumulative number of mismatches within all reported miRNA-mRNA interaction pairs.

##### Cumulative number of G:U pairs

The “GU_pair_plot.csv” file contains the cumulative number of G:U pairs within all reported miRNA-mRNA interaction pairs.

##### Cumulative number of adjacent mismatches

The “adj_mismatches_plot.csv” file contains the cumulative number of adjacent mismatches within all reported miRNA-mRNA interaction pairs.

#### Using R to produce plots

We provide a script to produce plots that can be used to manually infer targeting criteria based on your requirements. To use these, you need to have a valid version of R installed and correctly configured.

#### Clustered histogram of position specific properties and Minimum free energy ratio distribution

The script “create_plots.R” in the Workbench directory found at /ExeFiles/R-scripts/PAREameters, can be used to create plots such as the one below from the output of PAREameters.

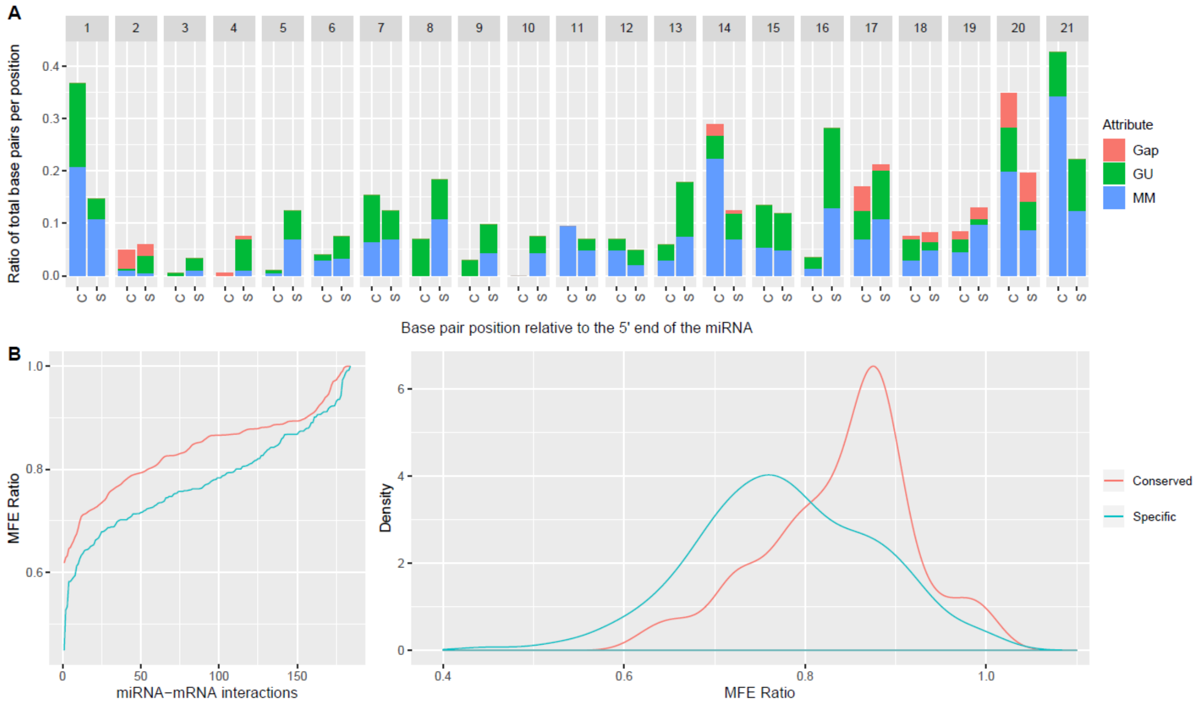


After the execution of the PAREamters pipeline, there will be two generated .csv files in the /ExeFiles/R-scripts/PAREameters directory. To create the plots, run the R script “create_plots.R” either through the command line or through your preferred R development environment. This will then produce you PDF files containing the plots. In addition, if multiple samples are included as input, statistical significance tests will also be performed.
